# MiSebastes: An eDNA metabarcoding primer set for rockfishes (genus *Sebastes*)

**DOI:** 10.1101/2020.10.29.360859

**Authors:** Markus A. Min, Paul H. Barber, Zachary Gold

## Abstract

Environmental DNA (eDNA) is a promising biomonitoring tool for marine ecosystems, but its effectiveness for North Pacific coastal fishes is limited by the inability of existing barcoding primers to differentiate among rockfishes in the genus *Sebastes.* Comprised of 110 commercially and ecologically important species, this recent radiation is exceptionally speciose, and exhibits high sequence similarity among species at standard barcoding loci. Here, we report new *Sebastes*-specific metabarcoding primers that target mitochondrial *cytochrome B.* Amongst the 110 *Sebastes* species, 85 unique barcodes (of which 62 are species-specific) were identified in our amplicon region based on available reference sequences. The majority of the remaining barcodes are shared by only two species. Importantly, MiSebastes yield unique barcodes for 28 of 44 commercially harvested species in California, a dramatic improvement compared to the widely employed MiFish *12S* primers which only recover one of 44. Tests of these primers in an aquarium mesocosm containing 16 rockfish species confirms the utility of these new primers for eDNA metabarcoding, providing an important biomonitoring tool for these key coastal marine fishes.

## Introduction

Traditional methods for monitoring marine fishes (e.g. trawling, visual surveys, seine netting) are costly and time and labor intensive, limiting the scope, scale, and reliability of assessments of ecologically and commercially important fish stocks (Murphy and Jenkins 2010). For example, the Channel Islands Marine Sanctuary employs visual monitoring by SCUBA divers to assess ecosystem health across 1,470 square miles of sanctuary. Despite great expense, they only survey 37 sites, recording just 110 of the 450-plus species of fish known to inhabit the coastal waters of Southern California—and only *once* per year. Given these limitations and the growing threat of climate change to coastal marine communities (Doney et al. 2011), new monitoring approaches that capture more diversity with greater frequency are required.

Environmental DNA (eDNA) has emerged as a potential new monitoring strategy for marine ecosystems (Kelly et al. 2014a; Thomsen and Willerslev 2015; Andruszkiewicz et al. 2019; Closek et al. 2019; Truelove et al. 2019). Environmental DNA is freely associated DNA shed by organisms into their environment (Taberlet et al. 2012). By isolating, amplifying, and sequencing eDNA, it is possible to identify species inhabiting a particular environment (Taberlet et al. 2012), providing a non-invasive and cost-effective alternative to traditional survey methods.

While early eDNA studies often focused on the detection of individual taxa using species-specific primers (e.g. Ficetola et al. 2008), recent efforts have focused on metabarcoding entire communities. eDNA metabarcoding allows detection of multiple species using a single set of “universal primers” that amplify a target barcode locus across a broad range of target taxa (Ji et al. 2013). When combined with next-generation DNA sequencing, eDNA metabarcoding can reconstruct community composition in many ecosystems (Bohmann et al. 2014), including marine ecosystems (Thomsen et al. 2016).

Metabarcoding of eDNA has been used to examine different marine communities along the Pacific Coast of North America (Port et al. 2016; O’Donnell et al. 2017), providing a potential alternative to the time-consuming and costly monitoring activities of local management agencies. However, central to successful monitoring of North Pacific coastal ecosystems is the ability to detect and distinguish among species within the genus *Sebastes*, commonly known as rockfishes. Rockfishes are ecologically important in North Pacific coastal food webs (Mills et al. 2007), and form the basis of a valuable commercial and recreational fishery (Lea et al. 1999; Love et al. 2002a; Williams et al. 2010). Of great concern, however, is that many of the commercially exploited species are in sharp decline. For example, landings of *Sebastes entomelas* and *S. flavidus* have declined by 79% and 62%, respectively, since 2000 (California Departmenmt of Fish and Wildlife 2020).

Despite these declines, monitoring *Sebastes* remains a significant challenge for managers. *Sebastes* includes approximately 110 species, at least 67 of which occur along the Pacific Coast of North America (Ingram and Kai 2014), where they occupy a wide range of habitats, from tide pools to depths greater than 1000 m (Love et al. 2002). Moreover, traditional survey methods can’t differentiate the early life history stages of rockfish species based on morphology (Thompson et al. 2016). While eDNA could be an effective alternative approach for monitoring *Sebastes*, existing universal teleost metabarcoding primers (e.g. the *12S* MiFish primers; Miya et al. 2015) cannot resolve *Sebastes* to the species level (Yamamoto et al. 2017). This shortcoming results from the rapid divergence of many species in this genus during the last million years (Hyde and Vetter 2007), resulting in rockfish species having identical or nearly identical DNA sequences at the mitochondrial *12S* locus targeted in the MiFish primers.

Effective monitoring of *Sebastes* will require new approaches like eDNA, but the application of eDNA hinges on identifying loci that evolve faster and have more sequence variability to distinguish among species. Here we present new metabarcoding primers that distinguish between species of *Sebastes*, providing an important new tool to advance eDNA monitoring of teleost fishes along Pacific coast of North America, and test these primers in a large marine mesocosm.

## Methodology

### 2.1 Primer design

We developed primers using *ecoPrimers* (Riaz et al. 2011), a software package that determines optimum barcode sites and their associated PCR primers based on a combination of metabarcode resolution and the conservation of the primer regions flanking the barcodes. First, we generated a database of input reference sequences, using the *in silico* PCR program *ecoPCR* (Ficetola et al. 2010) employing seven different primer sets *(control region, cytB, cox1, 16S, 12S, RAG2, ITS1*) from Hyde and Vetter (2007). We then imported the resulting sequences into *ecoPrimers*, specifying *Sebastes* as the target taxa and the order Perciformes as the non-target taxa. We specified a metabarcode length between 70 and 250 base pairs; a region shorter than 70 bp was unlikely to have the resolution necessary to distinguish between species, while a region longer than 250 bp is not well-suited to eDNA metabarcoding due to both DNA degradation in the environment and the short read length of next-generation sequencing platforms (Deiner et al. 2017; Jo et al. 2017). We then filtered the resulting list of primers based on the following criteria: (i) melting temperature (> 50 °C), (ii) both the forward and reverse primers targeting highly conserved regions unique to *Sebastes*, (iii) primer GC content (between 40% and 60%).

### 2.2. In silico primer validation

Following the methods of Riaz et al. (2011), we tested candidate primers *in silico* against the entire set of available DNA sequences in the EMBL-European Nucleotide Archive (release 133, standard sequences) to identify the number of target and non-target taxa amplified. Using the results from *in silico* PCR, we then examined the conservation and specificity of potential priming sites for our target and non-target taxa using DNA sequence logo plots (Crooks et al. 2004) and mismatch plots (Valentini et al. 2016) for both target and non-target taxa.

After identifying *Sebastes* specific primers, we utilized custom *R* (R Core Team 2013) functions from the *OBITools* package (Boyer et al. 2016) to determine the taxonomic resolution achieved by our different candidate metabarcodes. We then selected our top primer pair (hereafter called the “MiSebastes” primer set) based on its ability to distinguish the greatest amount of *Sebastes* taxa while minimizing non-target amplification. All code used for the generation of candidate primers and the *in silico* validation of the primers can be found at https://github.com/markusmin/MiSebastes.

### 2.3 In vitro validation of primers

To test the effectiveness of the MiSebastes primers in amplifying *Sebastes* DNA, we conducted PCR reactions using our primer set on DNA from 39 different species of rockfishes. We first extracted DNA from ethanol-preserved samples using the Qiagen DNEasy Blood and Tissue kit (Qiagen, Valencia, CA, USA), following the methods of Thompson et al. (2016). For each species, we had tissue samples from two different individuals, for a total of 78 samples. We amplified extracted DNA with a 25-μL reaction volume containing 1 μL of template DNA, 1 μL of both forward and reverse primer (10 μM), 12.5 μL of QIAGEN Multiplex Taq PCR 2x Master Mix, 2.5 μL of Q-solution, and 7 μL of nuclease-free water. Because the *Primer3Plus* software package (Untergasser et al. 2007) indicated the possibility of the MiSebastes primer forming hairpin turns, we included Qiagen Q-solution in PCR reactions to limit secondary structure formation and improve PCR results (Qiagen, 2010). Thermocycling parameters employed a touchdown PCR profile. After an initial 15-minute denaturation period at 95°C, the thermal cycle profile was as follows: denaturation at 94°C for 30 s, annealing at 64.5°C for 90 s, extension at 72°C for 30 s. These three steps were repeated 14 times, with the annealing temperature decreasing by 1.5°C each time, reaching a final ‘touchdown’ annealing temperature of 45°C. Following the ‘touchdown’ portion of the PCR, an additional 35 final cycles were carried out, with denaturation at 94°C for 30s, annealing at 60°C for 90 s, and extension at 72°C for 45s, followed by a final extension at 72°C for 10 minutes.

Following amplification, samples were Sanger sequenced by Laragen, Inc. (10601 Virginia Ave, Culver City, CA). We edited sequences and generated contigs of forward and reverse reads for 70 of the 78 samples (6 were poor quality reads and thus not used in further analyses and 2 samples did not amplify) using *Geneious* Version 2019.1 (Kearse et al. 2012). We then used the *BLAST* algorithm (Altschul et al. 1990) to compare sample DNA sequences to the NCBI reference database, recording the top three hits for each sequence, along with the percent identity, query cover, and accession number of the matching sequences (Supplementary Table 1). We aligned contigs from tissue DNA extractions in Geneious to detect intraspecific variation in our barcode and determine the variability of our barcode between species.

To visualize the DNA sequence similarity in our barcode locus across multiple *Sebastes* species, including the identification of species that share the same barcode, we created a distance-based phylogenetic tree. Briefly, we assembled all high-quality DNA MiSebastes metabarcode sequences into a single alignment, and then used the *phangorn* package in R (Schliep 2011) to generate a phylogeny based on hierarchical clustering using the unweighted pair group method with arithmetic mean (UPGMA).

### 2.4 In situ validation of primers

To test the utility of the MiSebastes primer set for eDNA metabarcoding, we collected water samples from the kelp forest tank at the California Science Center (700 Exposition Park Drive, Los Angeles, CA 90037), a 700,000 L aquarium exhibit containing 16 *Sebastes* species. We collected triplicate biological replicates of 1 L at both the surface and bottom of the well-mixed aquarium (total = 6 samples). We filtered water samples through a 0.2 μm Sterivex (MilliporeSigma, Burlington, MA, USA) filter using a gravity filtering method and stored filtered eDNA at −20°C for transport back to the lab at UCLA (Miya et al. 2016). We extracted eDNA from the filters using the Qiagen DNeasy Blood and Tissue kit (Qiagen, Valencia, CA, USA) following the modifications of Spens et al. (2017), in which ATL buffer and proteinase K are added directly to the sterivex filter cartridge. Cartridges were incubated overnight and then transferred to 1.7 mL tubes using a 3 mL syringe. Equal amounts of sample, AL buffer, and 0 °C ethanol were added. The protocol then followed the manufacturer’s instructions. We stored extracted eDNA at −20°C until PCR amplification.

We amplified extracted eDNA with the MiSebastes primers employing the same thermocycling parameters, above, but tripled the volume of template DNA from 1 μL to 3 μL to improve PCR sensitivity and performance. The PCR reaction volume was thus as follows: 3 μL of template DNA, 1 μL of each primer (10 mM) 12.5 μL of QIAGEN Multiplex Taq PCR 2x Master Mix, 2.5 μL of Q solution, and 5 μL of nuclease-free water. To minimize PCR dropout, we had three technical replicates for each sample by conducting all PCR reactions in triplicate (Doi et al. 2019), and all PCR reactions included negative and positive controls. Following PCR, we confirmed successful amplification and correct product size through electrophoresis on a 2% agarose gel.

To compare MiSebastes to MiFish primers (Miya et al. 2015), we repeated the above using the MiFish primers on the three surface samples. We used a similar touchdown PCR protocol, but the touchdown annealing temperatures began with an initial temperature of 69.5°C for 30 s and subsequently decreased by 1.5°C every cycle until 50°C was reached. The reaction volume was as follows: 1 μL of template DNA, 5 μL of each primer (2 mM), 12.5 μL of QIAGEN Multiplex Taq PCR 2x Master Mix, and 1.5 μL of nuclease-free water. Template volume for MiFish PCR differed from MiSebastes so that both yielded PCR products of similar strength.

We prepared PCR products for sequencing following the CaleDNA protocol (Meyer et al. 2019). The three technical replicates for each sample were sequenced separately for the MiSebastes primer set; the technical replicates for each sample were pooled before being sequenced for the MiFish primer set. Because MiFish and MiSebastes results were summed across all samples and technical replicates, this difference in methodology had a negligible effect on the results of the comparisons of overall primer performance.

We generated sequencing libraries through an indexing PCR using IDT for Illumina Nextera UD Indexes Sets A and D (Illumina, San Diego, CA, 92122) and KAPA HiFi HotStart Ready Mix (Kapa Biosystems, Wilmington, MA, USA). This second indexing PCR was performed using a 25 μL reaction mixture containing 12.5 μL of Kapa HiFi Hotstart Ready mix, 0.625 μL of primer i7, 0.625 μL of primer i5, and 10ng of template DNA, and used the following thermocycling parameters: denaturation at 95°C for 5 min, 5 cycles of denaturation at 98°C for 20 sec, annealing at 56°C for 30 sec, extension at 72°C for 3 min, followed by a final extension at 72°C for 5 min. We ran all indexed PCR products on a 2% agarose gels to ensure successful PCR and correct product size. We then cleaned the pooled samples using Serapure magnetic beads (Faircloth and Glenn 2014) and quantified cleaned PCR product concentrations using the high sensitivity Quant-iT™ dsDNA Assay Kit (Thermofisher Scientific, Waltham, MA, USA) on a Victor3 plate reader (Perkin Elmer Waltham, MA, USA). Lastly, we sequenced the libraries on a MiSeq PE 2×300bp at the Technology Center for Genomics & Bioinformatics (University of California – Los Angeles, CA, USA), using Reagent Kit V3 with 20% PhiX added to all sequencing runs.

We processed metabarcoding sequences from the MiSebastes primer set using the *Anacapa Toolkit* (Curd et al. 2019). To maximize identification of metabarcode sequences, we used *CRUX* to create a *Sebastes*-specific reference database following the standard *CRUX* parameters. As the *Anacapa Toolkit* assigns taxonomy to ASVs with a confidence score using a Bayesian Lowest Common Ancestor (BLCA) method, it can only assign one species to a particular ASV. As such, we developed an additional script to the *Anacapa Toolkit* to help assign taxonomy to sequences that are shared between multiple species for our metabarcode (https://github.com/markusmin/misebastes_annotator). This add-on for the *Anacapa Toolkit* identifies all species that share an ASV for our barcode and adds this information to the *Anacapa* taxonomy output file. We employed standard *Anacapa Toolkit* parameters and a Bayesian cutoff score of 40 to preserve the low-confidence annotations shared across multiple species for a given ASV to ensure the successful application of the MiSebastes-specific annotation script.

We processed metabarcoding sequences from the MiFish primer set using the *Anacapa Toolkit* (Curd et al. 2019) following standard parameters and a Bayesian cutoff score of 60 (Gao et al. 2017). We assigned taxonomy to MiFish data using FishCARD, a California Current-specific fish *12S*-specific reference barcode database (Gold et al. in review).

## Results

### 3.1 Design and in silico validation of primers

We generated 1,905 possible primer pairs distributed across five of seven target loci: 690 in *cytB*, 437 in *16S*, 430 in *CO1*, 341 in the control region, and 7 in *12S*; there were no candidate metabarcodes for *ITS1* and *RAG2.* However, after employing our primer design constraints, we identified only six suitable candidate primer pairs, all in *cytochrome B.* From these, we selected the primer pair with highest predicted resolution for *Sebastes*, herein referred to as the MiSebastes primer set (Table 1).

**Table 1.**
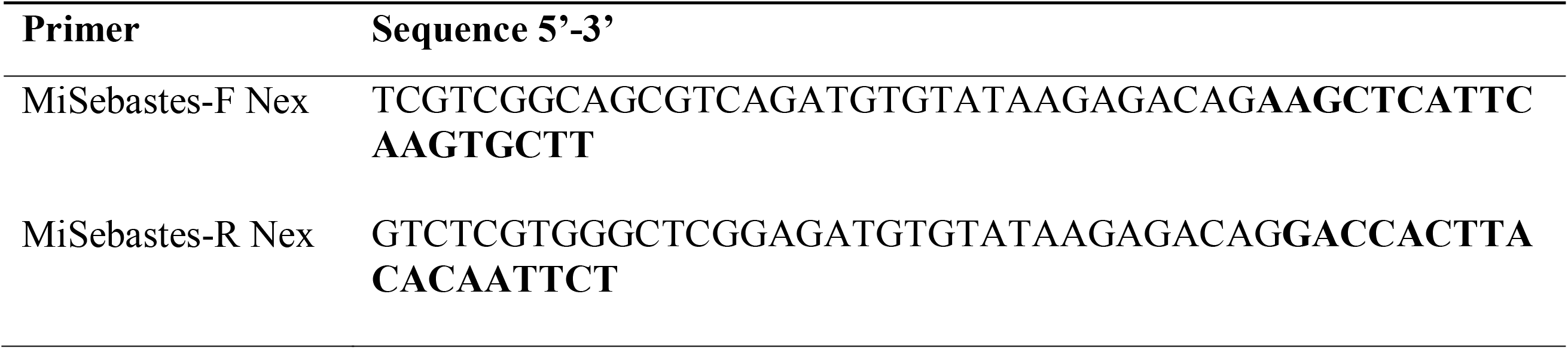
MiSebastes primer sequences (bold) and Nextera adapters.

Priming sites were highly conserved across the genus (Fig. 1a). For the forward MiSebastes primer, only 5 of the 100 species (*S. alutus, S. hubbsi, S. longispinis, S. oblongus*, and *S. scythropus*) have a single mismatch. However, the reverse priming region was less conserved; 49 species had zero mismatches, 41 species had one mismatch, five species (*S. hopkinsi, S. hubbsi, S. longispinis, S. pachycephalus*, and *S. variabilis*) had two mismatches, and one (*S. paucispinis*) has three mismatches. A complete species list with mismatches can be found in Supplemental Results Table 2.

**Fig. 1.**
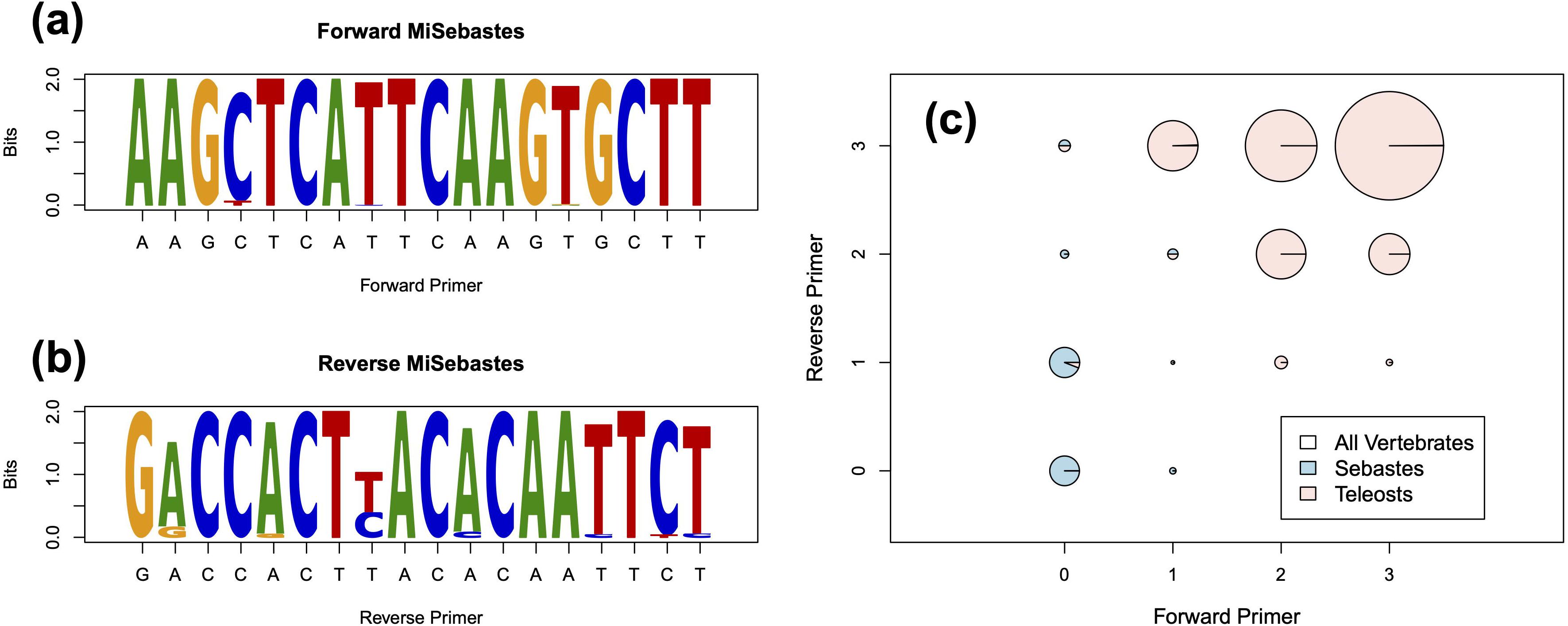
Three plots showing the conservation (a) and (b) and specificity (c) of the MiSebastes primer set for our target taxa (genus *Sebastes*) In (a) and (b), the size of the letter corresponds to the conservation of the forward (a) and reverse (b) primers at each base in the primer sequence. This indicates that the reverse primer is less conserved, especially for the eighth base. The mismatch plot (c) shows the number of mismatches in our priming regions among vertebrates, with the size of the circles corresponding to the number of taxa. (c) indicates that our priming regions are the most conserved for *Sebastes*, while other teleosts and vertebrates have much higher rates of mismatch.

MiSebastes amplified at least one unique sequence via *in silico* PCR for 62 of 100 species for which data is available in the EMBL-European Nucleotide Archive (release 133, standard sequences), and 43 of the 67 *Sebastes* species known to be found along the Pacific coast of North America. Owing to intraspecific variation in our targeted region, nine of the 62 species with unique barcodes had two unique sequences for our barcode. The remaining 38 species shared one of 13 unique sequences. Of these, 10 unique barcodes are shared by only a pair of species, one unique barcode is shared by three species, one unique barcode is shared by five species, and one unique barcode is shared by ten species (See Supplemental Results Table 2). It is unknown if the MiSebastes primer set is capable of amplifying and distinguishing ten *Sebastes* species (*S. cheni, S. diaconus, S. ijimae, S. itinus, S. nudus, S. simulans, S. varispinis, S. ventricosus, S. wakiyai*, and *S. zonatus)*, as there were no reference sequences available in the EMBL-European Nucleotide Archive (release 133, standard sequences) for the region targeted by our primers. Importantly, *in silico* PCR results revealed MiSebastes was highly specific to *Sebastes*, with all other vertebrates having at least three mismatches in either the forward or reverse primer and thus limiting annealing (Fig. 1c).

### 3.2 DNA barcoding and metabarcoding

MiSebastes successfully amplified DNA from 38 of 39 rockfish species. All amplified species had zero or one mismatch in the priming regions, except *S. hopkinsi*, which had two mismatches in the priming region. The only species that did not amplify was *S. paucispinis*, which had three mismatches in the annealing region of the reverse primer, with one of the three mismatches at the 3’ end.

Eight of the 38 amplified species exhibited intraspecific variation in the MiSebastes barcode region, with the remaining 30 species exactly matching the reference sequences in the EMBL database. Intraspecific variation was limited to one (*S. goodei, S. jordani, S. ruberrimus, S. wilsoni*) or two base pairs (*S. hopkinsi, S. mystinus, S. rastrelliger, S. rufinanus).* For our barcoded species, the DNA sequence similarity, intraspecific variation, and shared barcodes are shown in Fig. 2.

**Fig 2.**
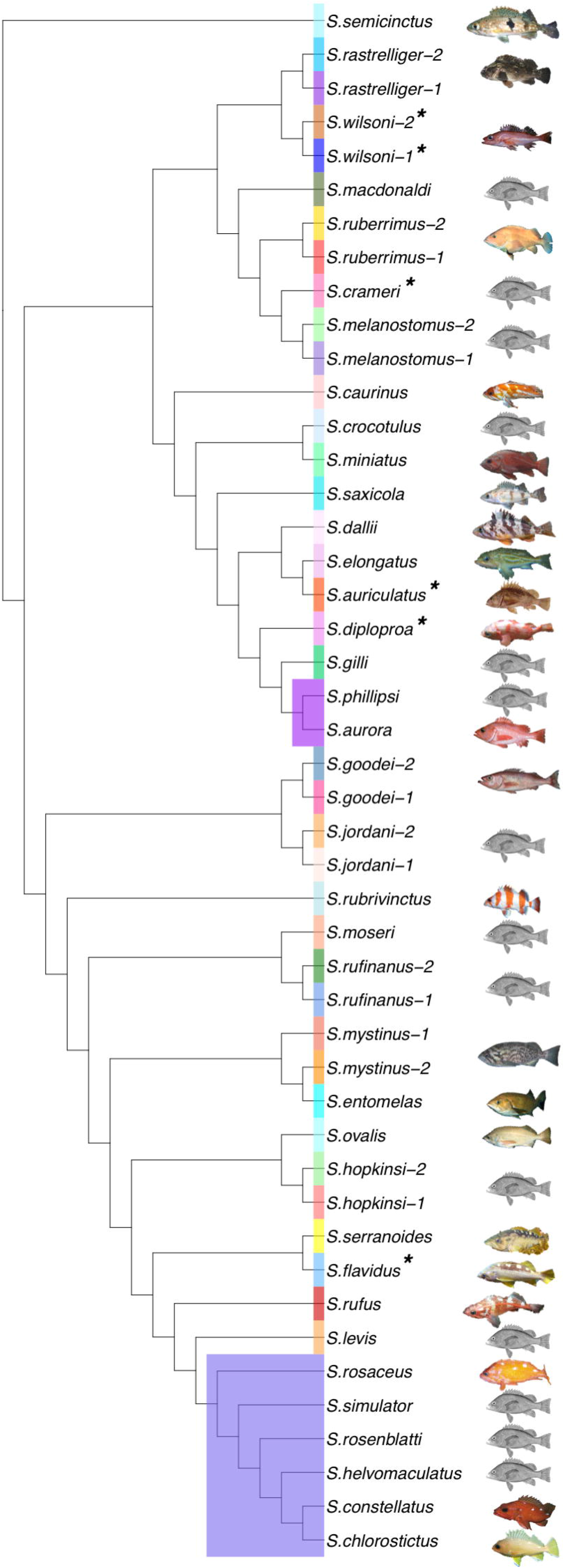
Single-gene phylogeny of MiSebastes barcode showing the similarity in our barcode across the 38 species that we successfully barcoded. Each color corresponds to a unique barcode. Species listed twice exhibit intraspecific variation in our barcode. Two barcodes were shared by more than one species, indicating the inability of MiSebastes barcode to discriminate species level identification for those taxa. An asterisk next to the species name indicates that this barcode in unique amongst the 38 species in the tree but not across the entire genus.

Sequencing of aquarium eDNA samples generated 11,064,598 sequences which passed filter from the MiSeq run on MiSebastes amplicons. Of these, 963,563 of these reads were assigned to ASVs through the *Anacapa Toolkit*, yielding an average of 160,594 (SE = 9650.81) sequences per sample. After filtering out only ASVs that were the correct length of 153 BP and constraining our output to select only ASVs assigned to taxonomy with a Bayesian cutoff score of 40, we detected a total of 34 ASVs for the MiSebastes sequences, assigned to one of 11 *Sebastes* species.

Of the 16 species of *Sebastes* known to be in the tank, we were able to detect and assign taxonomy to 11 using the MiSebastes primers. *Anacapa* assigned species-level taxonomy to 7 species (*S. caurinus, S. chrysomelas, S. rastrelliger, S. ruberrimus, S. rubrivinctus, S. serranoides*, and *S. serriceps*) with high (>70%) bootstrap confidence. An additional four species had a bootstrap confidence score of less than 50%. Three of these share a barcode with another species from *in silico* results; two of the species, *S. auriculatus* and *S. melanops*, share a barcode with a species not present in the aquarium, allowing positive identification of the eDNA sample to the species within the tank (Supplementary Table 2). The third species, *S. carnatus*, shares a barcode with *S. atrovirens*, allowing us to confirm that either or both were present in the eDNA sample but precluding distinguishing among them. The fourth species with a bootstrap confidence score of less than 50%, *S. umbrosus*, has a published reference sequence that is unique, but the ASVs we detected were shared with *S. lentignosus*, a species not present in the tank. There were five species in the tank not detected via eDNA: *S. atrovirens, S. constellatus, S. dallii, S. paucispinis*, and *S. mystinus.* We note that *S. atrovirens* shares a barcode with *S. carnatus*, and *S. constellatus* shares a barcode with nine other species. However, the other three species each have unique barcode sequences according to published reference sequences for the *cytB* gene.

Analyses from MiFish generated sequences yielded 20,615,456 sequences which passed filter from the MiSeq run on MiFish amplicons. Of these, 807,747 of these reads were assigned to ASVs through the *Anacapa Toolkit*, yielding an average of 269,249 (SE = 116,916.76) sequences per environmental sample. After processing sequences using the *Anacapa Toolkit*, MiFish recovered a total of 2,247 ASVs across the three samples. We constrained the output to select only ASVs assigned to taxonomy with a Bayesian cutoff score of 60, leading to the detection of 23 species amongst these ASVs. Notably, the MiFish primers were unable to distinguish any *Sebastes* spp. beyond the genus level except for *S. paucispinis* (Supplementary Table 3).

## Discussion

The Pacific Coast of North America is home to at least 67 species of *Sebastes*, (Ingram and Kai 2014) that are commercially and ecologically important, and many of which are in decline (California Deparment of Fish and Wildlife 2020). The MiSebastes primer set distinguished, *in silico*, 43 of 67 North American *Sebastes* species, and 62 of 100 known species. In contrast, with the exception of *S. paucispinis*, the MiFish primers (Miya et al. 2015) only yielded genus level identifications. Importantly, the MiSebastes primer set identified 28 of 44 commercially harvested species in California (California Department of Fish and Wildlife, 2020), and this utility was confirmed by distinguishing among many *Sebastes* in a large marine mesocosm. As such, the MiSebastes primers provide an important new metabarcoding tool that will greatly improve the teleost coverage of eDNA metabarcoding in North Pacific marine ecosystems, providing marine managers an important new monitoring tool.

### 4.1 MiSebastes *provides critical metabarcoding coverage for North Pacific coastal fishes*

Because different species of the genus *Sebastes* vary greatly in their functional roles (Love et al. 2002a), management agencies monitor and manage different species within this genus individually. As an example, *Sebastes* species account for 18 of the 178 monitored fishes by the Channel Islands National Park Service Kelp Forest Monitoring Program and 13 of the 35 monitored fishes by Reef Check California (Gillett et al. 2012). However, prior to this study, eDNA metabarcoding could not provide species-level resolution for this genus, severely limiting the application of this biomonitoring technique for North Pacific coastal ecosystems such as the Channel Islands. In contrast, MiSebastes primers can, in theory, distinguish among 11 of the 18 monitored rockfishes by the Channel Islands National Park Service Kelp Forest Monitoring Program and 7 of the 13 monitored rockfishes by Reef Check California.

While the targeted region of *cytochrome B* lacked sufficient variation to distinguish among all *Sebastes* species, the ability to distinguish 62% of *Sebastes* worldwide and 64% along the shores of North America is a substantial improvement over the MiFish primer set. As such, MiSebastes has the ability to substantially increase the effectiveness of eDNA metabarcoding in the Pacific by supplementing existing universal *12S* fish primers, maximizing the number of teleost detections in North Pacific coastal ecosystems.

### 4.2 eDNA results

The new MiSebastes primer set successfully recovered and identified 10 species of *Sebastes* from eDNA samples taken from an aquarium with 16 rockfish species. Not obtaining species-specific taxonomic assignments for four of these six unidentified species was expected based on *in silico* PCR results. The first two of these species, *S. atrovirens* and *S. carnatus*, share a barcode, and while we recovered that barcode in the metabarcoding data, we could not determine which of these two species it came from. The third undetected species, *S. constellatus*, shares a barcode with nine other species, making species-level identification using the MiSebastes primer impossible. The fourth undetected species, *S. paucispinis*, had three mismatches present in the reverse primer and failed to amplify with the MiSebastes primers, even from tissue samples.

It is unclear why two species known to be in the aquarium, *S. dallii* and *S. mystinus*, were not detected. While this result could be a limitation of the MiSebastes primers, it is more likely due to incomplete species inventory within the aquarium. Aquarium inventories at the California Science Center only recorded the individuals that were added to the tank—it does not record individuals that are removed due to mortality or transferred to other aquaria. Aquarium records show that only one individual of *S. dallii* and three individuals of *S. mystinus* were added to the tank (Supplementary Table 3), three years and seven years before water samples were taken, respectively. Neither species were visually observed during sampling, so it is possible that these two species were not in the tank during sampling, explaining their absence from our metabarcoding data. Another potential explanation may be that eDNA concentrations of these two species are too low to be detected, given the presence of only 1-3 individuals in a 700,000 L aquarium. Failure to detect rare species in aquaria using eDNA is common. For example, eDNA failed to detect a single ocean sunfish *(Mola mola*) from water samples of the Monterey Bay Aquarium’s Open Sea Tank (Kelly et al. 2014b).

Despite the MiSebastes primers failing to detect all *Sebastes* species within the aquarium, our results demonstrate a marked improvement over existing universal barcoding primers. The MiFish Universal Teleost primer set (Miya et al. 2015) is only able to provide species-level taxonomic resolution for one *Sebastes* species, a shortcoming noted by Yamamoto et al. (2017). Given the ubiquity and diversity of rockfishes along the Pacific coast of North America (Ingram and Kai 2014), the improved detection and identification of rockfish species using the MiSebastes primers represents a considerable advance in the applicability of environmental DNA to the marine ecosystems of this region.

An important feature of the MiSebastes primers is that they are highly specific to rockfishes, and do not amplify other teleosts. This feature allows the MiSebastes primer set to fill a critical gap in coverage in the MiFish primer set, much like the MiFish*-*elasmobranch (Miya et al. 2015) or MiBird (Ushio et al. 2018) primers. Ideally, a single metabarcode locus and primer set would allow the detection and identification of all rockfishes. However, our results strongly suggest that no single locus suitable for metabarcoding can provide complete species-level taxonomic resolution for *Sebastes*, a finding that is consistent with other metabarcoding studies that highlight taxonomic limitations of “universal” metabarcodes (Stat et al. 2017; Yamamoto et al. 2017; Valsecchi et al. 2020).

While our priming regions are highly conserved across the majority of species in the genus (no mismatches, n=52), at least 48 species have a mismatch in the annealing regions of the primers, with the location of the mismatch varying between species. Notably, some of the species in the aquarium had a known mismatch in the priming region, but their eDNA was still successfully amplified. However, mismatches in the priming region can lead to inefficient and sometimes unpredictable amplification (Klindworth et al. 2013) and thus may lead to higher rates of false negative detections for these species. Previous studies of other metabarcoding markers have found that just a single mismatch can lead to a 1000-fold underestimation of abundance (Bru et al. 2008), thus also limiting the use of this primer set to only presence/absence data for *Sebastes* species that have mismatches in the priming regions. These mismatch issues could potentially be overcome through the development of a degenerate version of the MiSebastes primers.

### 4.3 Intraspecific variation

Intraspecific variation can present a challenge to taxonomic assignment via metabarcodes, as there must be more variation between different species than different individuals for taxonomic assignment (Brown et al. 2015). Of the 38 *Sebastes* species from which tissue DNA was successfully amplified, 8 species exhibited intraspecific variation within the MiSebastes barcode. We found that for 7 of these 8 species both sequences had the highest *BLAST* alignment to the known species (Supplementary Table 1). However, for *S. mystinus*, intraspecific variation in the barcode led to one individual sequence having a better BLAST hit to *S. entomelas* despite originating from *S. mystinus* tissue. Furthermore, *in silico* PCR results showed intraspecific variation within ten species (*S. miniatus, S. oblongus, S. owstoni, S. rosaceus, S. saxicola, S. scythropus, S. steindachneri, S. taczanowskii, S. umbrosus*, and *S. vulpes).* Each have two different published sequences for the *cytochrome B* gene; for all species but *S. rosaceus*, both sequences are not shared with any other species in the genus and are thus unique to the species (Supplementary Table 2). For *S. rosaceus*, one sequence is unique, while the other is shared with nine other species. Additionally, multiple ASVs were detected for the same species from our aquarium samples. These results indicate that intraspecific variation within *the* MiSebastes barcoding locus may limit the effectiveness of this primer set for certain *Sebastes* species and highlight the difficulty of designing a single metabarcoding primer for this recently radiated clade of teleost fish.

### 4.4 Conclusion

Given the diversity of rockfishes throughout the northern Pacific Ocean (Ingram and Kai 2014), the detection of different species of this ecologically and commercially important genus with the MiSebastes primer set will greatly improve the utility of environmental DNA. Although the MiSebastes primers are unable to distinguish all species of the genus, it still represents a vast improvement for *Sebastes* over the currently available universal MiFish teleost primer set (Miya et al. 2015). Increased ability to identify *Sebastes* species from eDNA samples will provide tools to resource managers to help improve single stock management and aid in the recovery of species that are recovering after being significantly overfished, such as canary rockfish (*S. pinniger*) (Thorson and Wetzel 2015), Pacific ocean perch (*S. alutus*) (Wetzel et al. 2017), and cowcod (*S. levis*) (Dick and MacCall 2014). These three species are examples of the 28 commercially harvested rockfishes in California (of 44 total) that can be identified using the MiSebastes primer set. In addition, the MiSebastes primers can be used for gut content analyses of predators of rockfishes as well as for ichthyoplankton metabarcoding (Bucklin et al. 2016), providing additional insights into the ecology of the North Pacific and important information for fisheries management.

## Acknowledgements

We thank Emily Curd and Meixi Lin for their assistance with the primer design process. We thank Andrew Thompson and Dovi Kacev for providing us with DNA extracted from tissue samples. We thank Adam Wall for his help processing our Sanger sequencing data. We thank Keira Monuki, McKenzie Koch, Beverly Shih, Cristopher Ruano, and Nikita Sridhar for their assistance with lab help. Grants to M.A.M. from the UCLA Undergraduate Research Scholars Program and to P.H.B from the Howard Hughes Medical Institute supported this study.

---

**Supplementary Table 1** DNA sequences for Sanger sequencing of tissue samples barcoded with the MiSebastes primer set, along with the following BLAST results: top three hits for each sequence, percent identity, query cover, and accession number of the matching sequences.

**Supplementary Table 2** Amplicons for the region targeted by the MiSebastes primer set as predicted by *in silico* PCR results from ecoPCR. For each amplicon, the species that share the amplicon, the region where the species are found, and the number of mismatches in the priming regions are listed.

**Supplementary Table 3** Inventory of the species in the kelp forest aquarium at the California Science Center, along with detections by MiFish (using both the Fish CARD and CRUX databases) and the MiSebastes primer sets.

## Notes

### Competing Interest Statement

The authors have declared no competing interest.

